# Divergence of Functional Brain Network Configurations in Same and Other Race Face Recognition

**DOI:** 10.1101/2025.02.06.636962

**Authors:** Jessica L. Yaros, Bethany A. Houalla, Liv McMillan, Myra S. Larson, Guanqiao Yu, Martina Hollearn, Blake A. Miranda, Robert Jirsaraie, Diana A. Salama, Paul E. Rapp, Michael A. Yassa

## Abstract

The “Other-Race Effect” (ORE) refers to the enhanced memory individuals have for faces within their own racial group compared to other races. This effect is attributed to limited exposure to different races and social-motivational factors affecting face processing. While past research has explored this effect through neuroimaging methods, the precise neural mechanisms that underlie the ORE remain a subject of ongoing investigation. Previous studies have largely adopted a modular perspective, concentrating on the differential activation of face-processing regions when comparing same-race and other-race facial recognition behaviors. However, given the multifaceted nature of the ORE, an exclusive focus on specialized regions may miss the pivotal role played by non-face-preferential brain areas in modulating this phenomenon. In the present study, we take a broader data-driven perspective using graph-theoretical techniques to investigate whole-brain network disparities in same- and other-race facial recognition. Our findings reveal a substantial impact of race on functional connectivity during face memorization. While other-race face recognition benefitted from higher local efficiency during encoding, reduced local efficiency was advantageous for memorization of same-race faces. Notably, successful other-race recognition was disproportionately supported by locally efficient processing in regions among the ventral and dorsal attention networks.

## Introduction

Faces are a category of visual objects that we interact with constantly from the moment of birth. This near ubiquitous exposure to faces during development tunes systems in the brain to expertly process facial information^1,2^. Entire job families revolve around the ability to process facial information; personnel at airports, banks, and government facilities must routinely match the faces of people to their photographs. And yet, research demonstrates people struggle with this task even when identifying faces of their own race group ^3,4^. It should be no surprise, then, that facial recognition for other-race faces can be even more challenging. Research has shown this disparity is due to a lack of regular interaction with members of these groups from a young age, as well as socio-motivational differences in processing of faces implicitly labeled as ‘out-group’^5,6^.

This so-called ‘Other-Race Effect (ORE)’ has serious ramifications in criminal justice system proceedings. In the United States a disproportionally large percentage of wrongful convictions discovered to date involve mistaken cross-race facial identifications^7,8^. Despite these numbers, few policies specifically combating the ORE have been implemented by law enforcement agencies or the court system^9^. A better understanding of the ORE’s neural basis can therefore help bolster the case that new policies should be put in place to mitigate negative consequences of the intersection of race and memory on eyewitness testimony.

Studies have shown the ORE involves differential engagement of specialized facial processing patches across the brain when participants view faces of their own race versus other races. The first-ever MRI study of the ORE found that participants passively viewing faces had higher activity in the right Fusiform Face Area (rFFA) to same-race (SR) faces, while the magnitude of the lFFA signal was correlated with an advantage in subsequent memory for SR relative to other-race (OR) faces ^10^. Over the years studies have continued to find variations between SR/OR face activity in the FFA and other cortical face patches like the Occipital Face Area (OFA)^11–13^.

While behavioral research has indicated a role for differential attentional processing of SR and OR faces in the ORE^1,5,14,15^, there is a dearth of research investigating brain regions associated with attention, or, for that matter, brain regions beyond those exhibiting face-preferential activity. One particular study highlights the importance of shifting focus beyond face-selective regions^16^: Reduced activity in cognitive control networks during face encoding was associated with failure to recognize OR but not SR faces. Furthermore, functional connectivity between the FFA and Dorsal Attention networks was more predictive of success for SR relative to OR recognition. These results demonstrate not only the importance of attentional and executive control regions during encoding of faces, but also a disparity between how SR and OR faces are processed beyond the core face patches in the brain. Overall, these results indicate that regions need not execute face-preferential activity to play a mediating role in the ORE.

The study^16^ demonstrates that a focus on the magnitude of activity in face-selective regions may miss crucial information on how disparate regions of the brain interact with one another. The ORE, which is behaviorally modulated by both past expertise and memory, visual experience, and attentional and motivational processes, is complicated and arises from a correspondingly complex interplay of brain networks. Research in the field of network neuroscience demonstrates the value of taking a holistic whole-brain approach to studying cognitive function by analyzing the brain as a graphical system of nodes (or brain regions) connected by edges (or functional connectivity estimates)^17–19^. A graph-theoretic approach allows neuroscientists to analyze the topology of brain interactions, and how brain network configurations are altered by changing cognitive task demands. One common analytic approach is to calculate the ease with which information may propagate between brain regions, where efficiency in communication between a pair of nodes is inversely proportional to the shortest path length between those nodes^20^; i.e., the shorter the path (or number of steps) between two nodes, the greater the efficiency of information exchange. This rule can be applied to characterize how well brain regions are integrated with one another (global efficiency), but also how fault-tolerant a network’s nodes are to perturbation (local efficiency/segregation).

One study demonstrating this interplay between global and local efficiency found that functional brain networks exhibited high local efficiency at rest, but during a working memory task, connectivity shifted to a high globally efficient configuration^21^. Furthermore, across participants, increasing global efficiency was correlated with improvements in working memory. A separate study using an n-back test found better working memory performance is associated with decreased local efficiency across young and older populations^22^, and for the young experimental group particularly, global efficiency predicted working memory capacity. These investigations suggest that more cognitively demanding tasks may rely on greater information integration across the brain via a reduction of local efficiency and facilitation of global efficiency. However, there is seemingly conflicting evidence that improved performance on difficult tasks over time is associated with increased modularity (which is related to increased local efficiency). One study found that after six weeks of training on a working memory task, performance and modularity were positively correlated, suggesting a relationship between task automation and modularized locally efficient processing^23^.

Reports like these show that the brain dynamically reconfigures large-scale functional connections to meet the demands of changing contexts. However, to our knowledge, no study has tested graph network reconfigurations on two contexts as closely related as same- and other-race face recognition. One might assume that no difference would be found in widespread network processes supporting SR versus OR faces. However as discussed, memory performance is associated with brain topological changes^21–23^ and the ORE—a memory deficit for OR faces—involves differential engagement of regions across the brain. In the present study we test whether the ORE is associated with significant differences in the global and local efficiency with which whole-brain networks functionally organize during encoding and retrieval of SR and OR faces.

## Methods

### Participants

This study protocol was approved by the Institutional Review Board (IRB) at the University of California, Irvine, and complies with IRB guidelines and regulations. Participants were screened for eligibility through a secure online questionnaire on REDCap^24^. Strict inclusionary criteria required participants to be right-handed with normal corrected vision, self-identify as East Asian or Southeast Asian, have no MRI contraindications such as metal implants, and no major neurological, psychiatric, or substance-use conditions. 27 recruited subjects (14 Female, 13 Male; mean age 19.93; SD 1.36; age range 18-22) provided written informed consent and were compensated for their participation. During the study, the participants filled out a series of questionnaires before and after scanning, as well as performed facial recognition tasks in the scanner. Because awareness of the true nature of the study could bias performance^25,26^, participants were only informed that they would be administered a facial recognition task, with no mention of the race component of the study. After the scan they were debriefed about the full study purpose and consented once again.

Of the initial sample, five participants were excluded from the analysis. Reasons for exclusion included participants not meeting final inclusionary criteria, demonstrating chance performance, missing 20% or more of trials, as well as technical difficulties with the scanner. This yielded a final sample of 22 subjects (11 Female, 11 Male; mean age 19.52, SD 1.29; age range 18-22). Of the final participants, 9 identified as East Asian and 13 as Southeast Asian. Two of these subjects only completed three of the four scan runs.

### Mnemonic Discrimination Task

This experiment was designed to test participants’ retrieval memory for faces of their own and another race. The task was adapted from our prior work^18^ for use in the MRI scanner ^27^. Research from unpublished pilot studies found that Asian participants displayed greater memory deficits for Black faces than White faces. We therefore limited our design to include only Asian and Black faces to focus on the larger behavioral differential in our sample. Since all subjects identified as Asian, for the purposes of this manuscript we refer to Asian faces as ‘Same-Race’ (SR) and Black faces as ‘Other-Race’ (OR).

### Stimulus Set

This experiment used the same stimulus set developed for our prior study^27^. All faces were generated using FaceGen Modeller 3.5. Asian and Black faces were created using the ‘Generate’ tool within sub-groups for ‘Asian’ and ‘African’ racial origins. Half of the faces were randomly selected as ‘parent’ stimuli to serve as templates for lure distractors. These lures were created by running the Genetic Randomness algorithm on the parent faces, to apply normally distributed perturbations with means proportional to inputted values of 20%, 30%, 40%, and 50%; this introduced variation in how similar lure faces were to ‘parent’ faces. The current study groups all similarity levels together to increase trial counts for subsequent neuroimaging analysis. However, refer to our earlier study ^27^ for an analysis of the interaction of facial race and perceptual perturbations. Our results demonstrated that despite controlling perceptual differences between memorized and subsequent distractor faces, participants performed more poorly on all but the most distinct OR faces. These results were not found in a control match-to-sample working memory task, indicating that the perceptual differences are exacerbated when longer term memory is required.

### Task Design

This task, programmed in PsychoPy v1.85.2 ^28^, was structured as an event-related design with 8 interleaved study and test phases. In Study Phases, participants were instructed to memorize each of 22 consecutively presented faces. They were simultaneously expected to indicate via button press whether faces were shifted to the left or right of the screen, to ensure they were attending to the task. In each Test Phase participants were again shown a series of 22 consecutive faces, half of which were directly repeated (target repeats) from the Study Phase just prior (Figure 1A). The other half of the faces were new (lure distractors). For each Test trial participants were asked to identify via button press whether the face was an exact repeat from the preceding Study Phase (‘Same’/’Old’) or whether they had never seen it before (‘New’/’Different’). A response of ‘same’ to a repeated face indicated successful recognition (i.e., target hit) while a response of ‘different’ to a new face indicated successful mnemonic discrimination (i.e., lure correct rejection). Across both Study and Test Phases, face stimuli were pseudorandomized and evenly divided amongst race and gender categories. Each stimulus was presented for 3.0s with a 1.5s intertrial interval (ITI).

**Figure 1:**
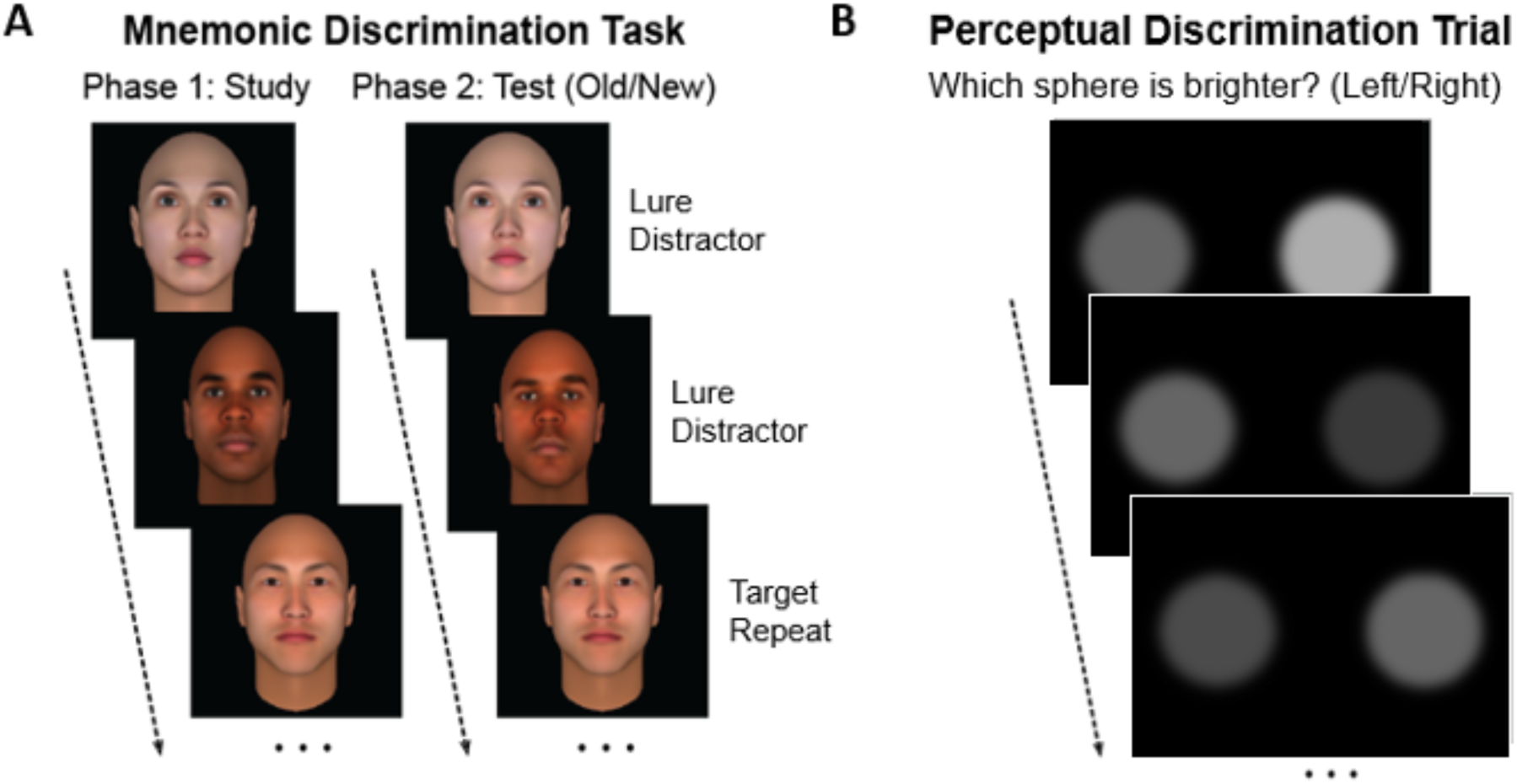
Mnemonic Discrimination and Perceptual Discrimination Task Designs: (A) The Mnemonic Discrimination task contained 8 blocks, with each block containing a Study and Test Phase. During the Study Phase participants memorized consecutively presented faces. They simultaneously indicated whether faces were shifted to the Left or the Right of the screen’s center (not displayed). During the immediately following Test Phase they were asked if each of the presented faces were shown during the Study phase. Half of the faces were shown during study (target repeats), and the other half were new (lure distractors). (B) 3 perceptual discrimination trials were randomly displayed during each phase of the Mnemonic Discrimination task. In these trials participants indicated which of two gaussian blurred spheres were brighter (Left or Right). If correct, the brightness of the spheres became more similar, and thus more difficult to discriminate. If incorrect, the spheres would become more distinct in brightness, and hence more discriminable. Participants performed these discriminations for the entire trial duration. For all trial types, stimulus duration was 3.0 s and the intertrial Interval (ITI) was 1.5s.

In addition to trials of interest, the task also employed perceptual discrimination trials to serve as an implicit baseline when subsequently modeling the fMRI BOLD response (Figure 1B)^29^. The perceptual discrimination trials required subjects to indicate which of two gaussian-blurred circles appeared ‘brighter’. The trials were adaptive, adjusting based on participant response during a 3-second trial duration. When answered correctly, the circles would become more similar in luminosity, and hence more difficult to discriminate. When incorrect, the circles would diverge in luminosity. Three perceptual baseline trials were pseudorandomly presented per phase.

### MRI Data Acquisition

Neuroimaging data were acquired on a 3.0 Tesla Siemens MAGNETOM Prisma scanner, using a 32-channel head coil at the Facility for Imaging and Brain Research (FIBRE), part of the Campus Center for Neuroimaging (CCNI) at the University of California, Irvine. A high-resolution three-dimensional (3D) magnetization-prepared rapid-gradient echo (MP-RAGE) structural scan was acquired at the beginning of the session, with the following parameters: 2300 ms repetition time (TR), 2.38 ms echo time (TE), 240 slices, 0.8 mm isotropic resolution, 256 mm field of view (FoV), and 8-degree flip angle. Parallel acquisition was conducted in the GRAPPA mode with reference line phase encoding (PE) of 24 and an acceleration factor of 3. Each of four functional MRI scans were acquired using a multiband echo-planar imaging (EPI) sequence with the following parameters: TR = 1500 ms, TE = 34 ms, 64 slices, 2.1 mm isotropic resolution, 202 mm FoV, 75-degree flip angle, and multiband acceleration factor of 8. Visual stimuli were presented on a BOLDScreen32 LCD monitor mounted onto the back of the bore. Participants viewed the monitor via a single mirror attached to the head coil.

During the session, structural scans were collected first and used to align all subsequent scans to the anterior commissure-posterior commissure line (AC-PC line). Following that, two blocks of study and test phases were presented during each functional run. Half-way between the four runs, subjects were given a break from the task during collection of a resting state scan.

### Preprocessing and Denoising

Structural and functional scans were preprocessed and denoised using the CONN Toolbox^30^. Functional scans were realigned and unwarped, centered, slice-time corrected, flagged for outliers, segmented, aligned to MNI space, and smoothed using a spatial convolution Gaussian kernel of 4 mm full width half maximum (FWHM). Structural scans were skull-stripped, segmented, and aligned to MNI space. A denoising pipeline was implemented on each scan run to create regressors for potential confounding effects. The nuisance regressors included the following: CSF and white matter noise components using anatomical component-based noise correction (aCompCor)^31^, parameters to minimize motion-related variability via 6 realignment regressors and their first-order derivatives, and scrubbing/censoring covariates using ART-based flagging of outlier volumes with framewise displacement above 0.9 mm or global signal change above 5 standard deviations^32^. Additionally, regressors were created for the experimental task effects by convolving stimulus onset and duration with a canonical hemodynamic response function (HRF). These included regressors for each of 16 conditions of interest corresponding to the unique combinations of race (SR or OR), phase (Study or Test), trial type (Target Pair or Lure Pair) and accuracy (Correct or Incorrect), as well as task effects of no interest (non-response trials, and task instruction reminders). Perceptual discrimination trials were not explicitly modeled, serving as an implicit baseline against which to compare increases or decreases in connectivity during the event types of interest^29^. Lastly, a linear component was added to model scanner drift. After regression, high-frequency information was preserved using a high-pass temporal filter [0.008 inf].

Quality control checks were run after preprocessing and denoising was complete. We found that across all subjects and runs, motion and global signal change were minimal. Mean framewise displacement was .13 mm with a standard deviation of .04 and mean global signal change was .83 with a standard deviation of .03. Because two subjects were missing one run due to scanner glitches, they were outliers for quantity of included volumes. However, both subjects maintained enough trials to safely model each condition (means of 16.19 and 15.89 and minimums of 9 and 12 trials per condition, respectively). Additionally, using the IQR outlier detection method, two subjects were established as potential outliers for mean motion (.23 and .24 mm). Given that these are still relatively low mean framewise displacements, we chose to include all data from these subjects as well.

### Whole-brain Generalized Psycho-Physiological Interaction Analysis

To derive whole-brain task-modulated connectivity data for subsequent graph theoretical analysis, we ran a generalized Psycho-Physiological Interaction Analysis (gPPI)^33^ on seed and target regions of interest (ROIs) across the entire brain^34,35^. The brain was partitioned into 246 ROIs using the Human Brainnetome Atlas parcellation scheme (Figure 2) ^36^. The fully preprocessed and denoised timeseries for each voxel were concatenated across runs, and then averaged within each of the ROIs. Next a general linear model (GLM) was fit for each ROI including as predictors: a) the HRF-convolved main task effects (i.e., the psychological factors), b) the timeseries for the seed ROI (i.e., the physiological factor), and c) the product of the psychological factors and the physiological factor (i.e., the interaction term). Modeling task-dependent connectivity for all pairs of ROIs required constructing 245 GLMs for each of 246 target ROI, for a total of 60,270 separate regressions. PPI terms for the 16 conditions of interest were then assembled into separate 246x246 connectivity matrices.

**Figure 2:**
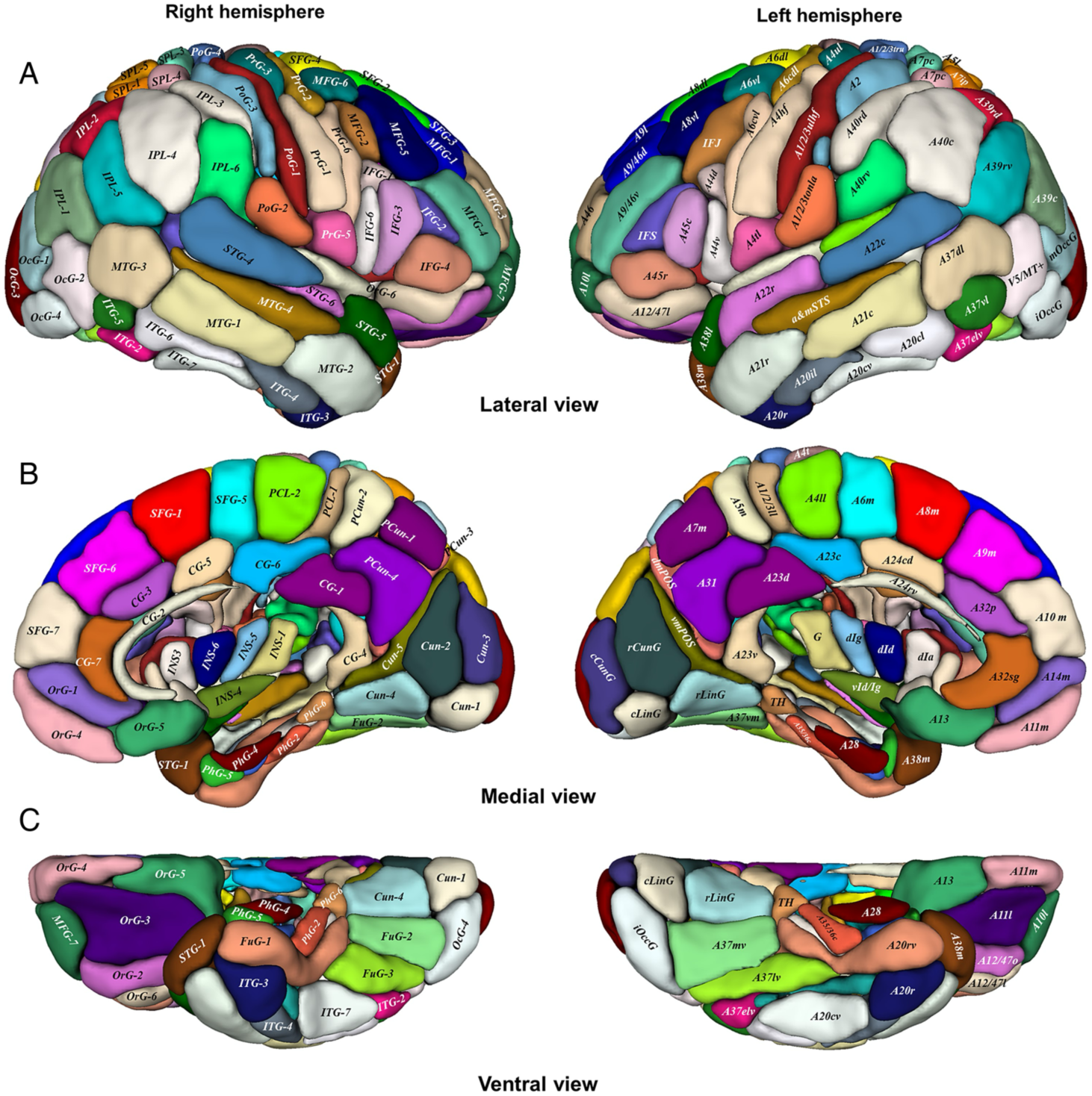
Parcellation scheme from the Brainnetome Atlas. All 246 regions were used for analysis of connectivity across the entire brain. (A) lateral view, (B) medial view, (C) ventral view. Figure reused from Fan et al., “The Human Brainnetome Atlas: A New Brain Atlas Based on Connectional Architecture,” Cerebral Cortex, 2016 under the terms of the Creative Commons Attribution Non-Commercial License (http://creativecommons.org/licenses/by-nc/4.0/).

Because the estimated effects of a seed ROI on a target ROI are not necessarily equal to the effects of the reverse couplings, gPPI functional connectivity matrices are not symmetric. This quality distinguishes gPPI models from the traditional bivariate-correlation weighted GLMs used in resting state connectivity analysis. This lack of symmetry cannot be used to infer directionality of information flow between source and target ROIs, or whether coupling is direct or mediated by other ROIs^37^. Because there is no consensus on how to interpret this directedness in gPPI estimates, we symmetrized all connectivity matrices by averaging the estimates across the matrix diagonal^38,39^.

### Graph Representation

In graph theory, a graph is a mathematical structure that models pairwise relations between objects. Graphs (or networks) are defined as sets of nodes connected by edges. In the context of network neuroscience, we can model brain regions as nodes and functional associations between regions as edges^40^.

Graphs are represented by adjacency matrices, which are square matrices with quantities of rows and columns equal to the number of (in this case) 246 brain regions (Figure 3). The value of elements in a matrix corresponds to the associations between each pair of brain ROIs. Here, our associations are the gPPI interactions between brain regions during task conditions. Because those interactions are averaged across the diagonal to be symmetric, these networks are considered undirected, such that only one edge is possible between any pair of ROIs.

**Figure 3:**
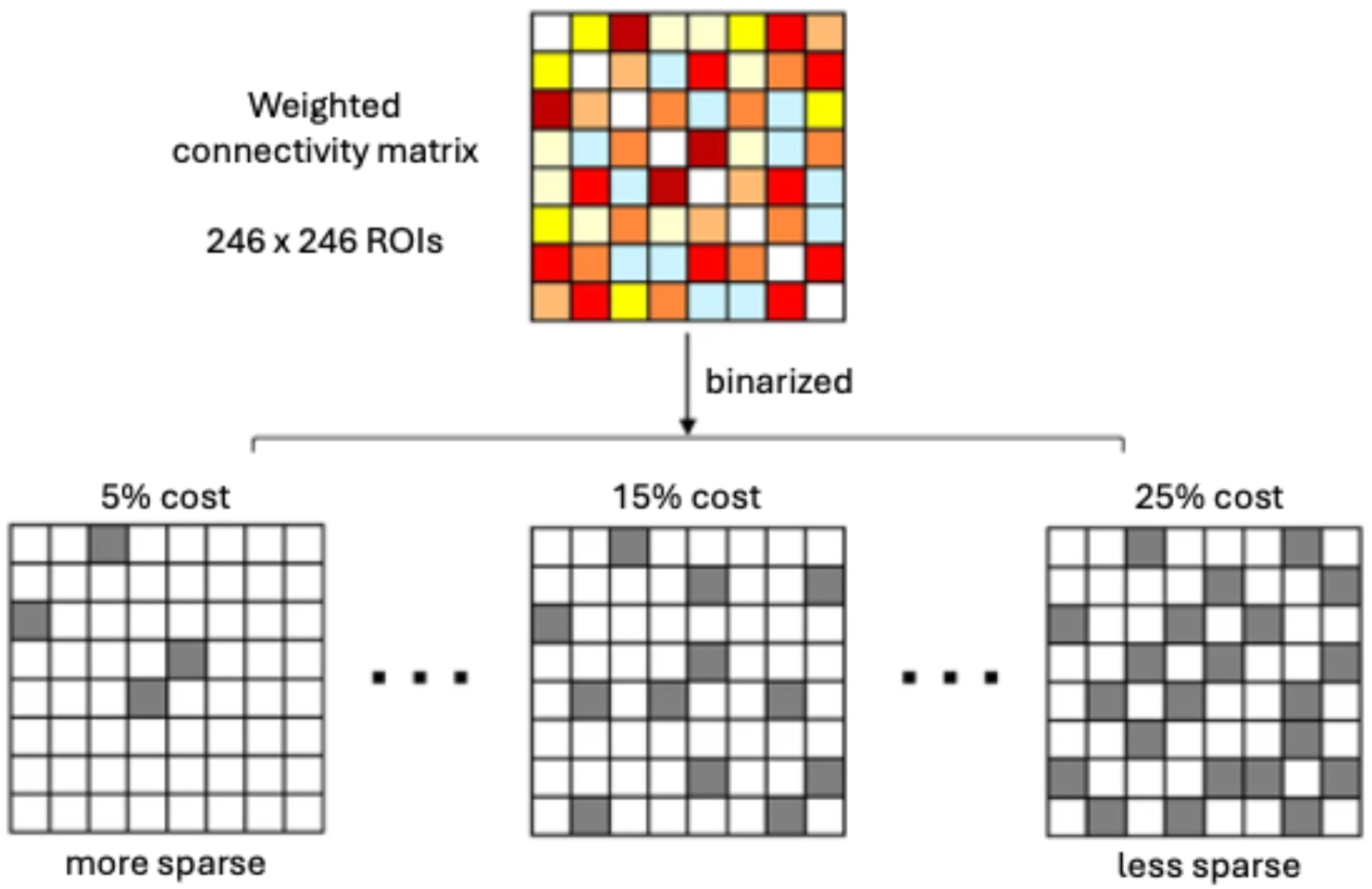
Graph Representations of whole-brain functional connectivity. Pairwise gPPI functional connectivity estimates were assembled into 246 × 246 connectivity matrices for each unique participant/task combination. Correlation matrices were then binarized at 5 cost levels, ranging from 5% cost (keeping the 5% strongest connections, most sparse) to 25% cost (keeping the 25% strongest connections, less sparse) using 5% step sizes. Figure modified from “Adams et al., “Functional network structure supports resilience to memory deficits in cognitively normal older adults with amyloid-β pathology,” Scientific Reports 2023 under the terms of the Creative Commons Attribution 4.0 International License (http://creativecommons.org/licenses/by/4.0/).

In network analysis, adjacency matrices are commonly binarized to contain only values of 1 and 0, indicating whether an association (or edge) exists (Figure 3). This requires an a priori decision regarding what constitutes a meaningful, or strong enough connection strength to be represented within a network. Rather than choosing one cut-off connection strength, five distinct thresholds ranging from .05 - .25 were applied to each individual connectivity matrix ^21,41–43^. Therefore, for each subject, each of the 16 connectivity matrices were described by five binarized adjacency matrices representing the strongest 5%, 10%, 15%, 20%, and 25% of functional interactions in the network. This range of costs was selected because the complexity and non-random topology of the brain is most observable at lower connection densities (number of edges per all possible edges)^42^. This allowed us to calculate graph metrics (described in the next section), while ensuring that the number of edges per graph was consistent and comparable across conditions and subjects^43^. Once metrics were calculated, they were averaged across costs to derive single measures per condition^21^. Matrix manipulation and analysis was all performed in MATLAB R2020A^44^.

### Graph Metrics

Graph theoretical metrics were calculated using the Brain Connectivity Toolbox^45^ (brain-connectivity-toolbox.net), a MATLAB toolbox for network analysis of structural and functional brain connectivity data. Our analysis focused on characterizing global and local efficiency^20^ of functional networks during the task conditions.

In graph theory, the efficiency of a pair of nodes is defined as the multiplicative inverse of the shortest path between that pair^20^. The global efficiency of each brain network was calculated by taking the average efficiency of all combinations of node pairs. This computation is formalized in Equation 1:

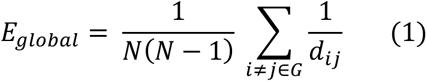

where *N* is the number of nodes in graph G; 1/*d_ij_* is the inverse shortest path length between a given pair of nodes *i* and *j, i*≠*j*∈G states the sum is taken over all pairs of distinct nodes in graph *G.* In contrast, the mean local efficiency of individual nodes was calculated by computing the efficiency of each nodal subgraph:

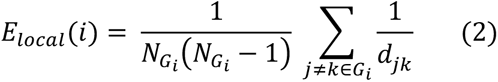

where *G_i_*, is the subgraph of node *i*; *N_Gi_* is the number of nodes in *G_i_* (i.e. the direct neighbors of node *i)*; 1/*d_jk_* is the inverse shortest path length between two nodes *j* and *k; and j* ≠ *k* ∈ *G_i_* indicates that the summation is over all pairs of distinct neighbors of *i* within the subgraph. To convert this to a network-wide metric, we compute the mean local efficiency of the entire graph by averaging the local efficiencies of each subgraph in graph G:

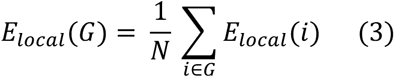

where N is the number of nodes in graph G; *E_local_* (*i*) is the local efficiency of node i as defined in equation 2; and *i* ∈ *G* states the summation is over each node’s subgraph. From here on, ‘local efficiency’ refers to that of the graph (*E_local_(G))*, rather than individual nodes.

While both metrics characterize efficiency of information exchange, global efficiency reflects how well integrated a network is, while local efficiency is a segregation metric that reflects the fault tolerance of a network. In general, systems with efficient information exchange tend to demonstrate both high global and local efficiency. Figure 4 depicts how connectivity within graphs can result in low versus high efficiency configurations.

**Figure 4:**
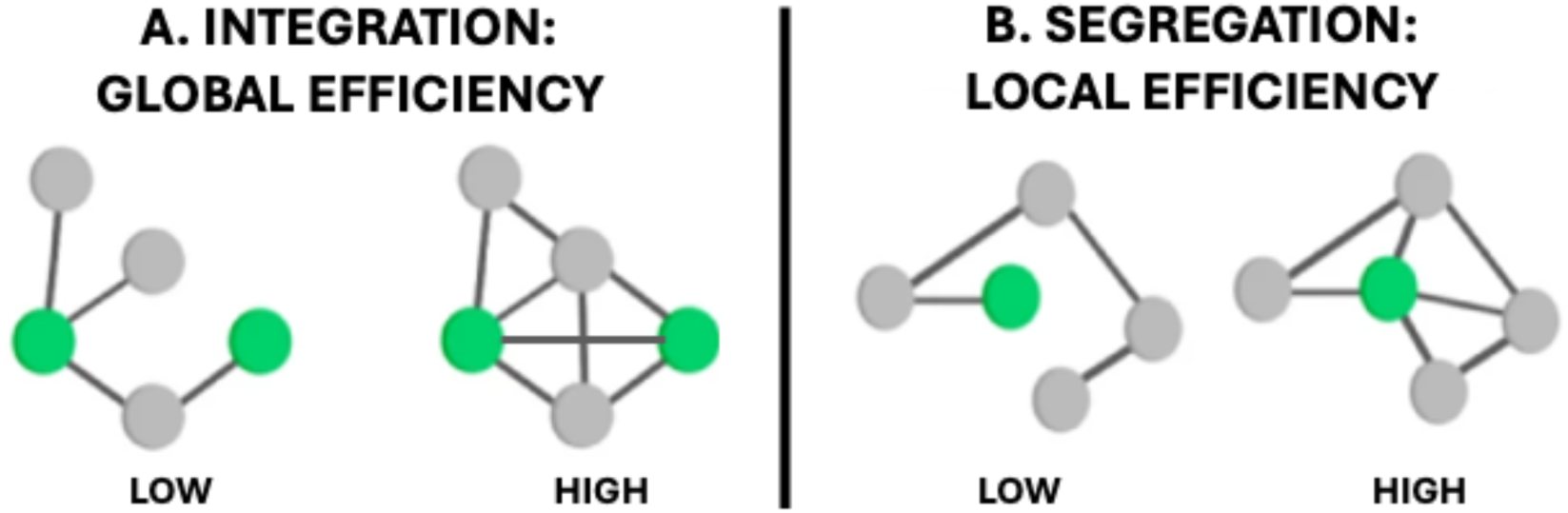
Global and network-wide local efficiency metrics were computed for each task-based functional brain network. Nodal efficiency is defined as the multiplicative inverse of the shortest path between a pair of nodes. A) Global efficiency is the average of the nodal efficiency across all node pairs (Eq. 1). A highly connected graph with short paths connecting disparate nodes will results in a more integrated, globally efficient network. B) To compute mean local efficiency, the mean efficiency of each nodes’ subgraph is computed (Eq 2). Next the subgraph efficiencies are averaged for a network wide local efficiency metric (Eq. 3). Local efficiency is a segregation metric that captures the fault tolerance of a network. I.e., how similarly the network can perform if a node is removed). A graph that contains nodes with large subgraphs will demonstrate high mean local efficiency and is considered robust. Well-connected systems tend to demonstrate both high local and global efficiencies. Figure modified from Borne et al., “Unveiling the cognitive network organization through cognitive performance.,” Scientific Reports, 2024 under the terms of the Creative Commons Attribution 4.0 International License (https://creativecommons.org/licenses/by/4.0/).

## Results

### Behavior

There are several common behavioral indices of the Other-Race Effect. These include greater target hits to same-race faces, greater false alarms to other faces, greater sensitivity to same-race faces, and more liberal response criterion to other-race faces. Because we expected effects to take on these directions, we ran 1-tailed paired t-tests. Participants accurately identified faces as repeated (target hits) for same-race (SR) and other-race (OR) faces with the following mean proportions: [SR: x̄ = .552, σ = .111; OR: x̄ = 0.577, σ = .109]. Subjects incorrectly recognized lure distractors as repeated (lure false alarm) with the following proportions: [SR: x̄ = .485, σ = .097; OR: x̄ = 0.536, σ = .093]. While it was not expected that the mean proportions for target hits would be larger for OR than SR faces, lure false alarm ratios followed the anticipated pattern. The proportions were used to calculate metrics from signal detection theory – both sensitivity, *d’* (Equation 4) and response criterion, c (equation 5):

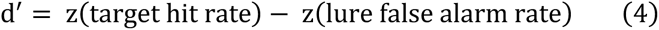

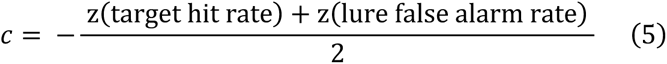

The Sensitivity, *d*’, corrects for the tendency to respond “old/same” inaccurately, providing a measure of participants’ ability to discriminate between old repeated and new distractor faces. The mean *d’* for the subjects were [SR: x̄ = 0.1726, σ = 0.183; OR: x̄ = 0.1047, σ = 0.2253] (Figure 5A). A paired t-test finds that same-race discriminability trends slightly larger than OR discriminability [t(21) = 1.542, p = 0.069, r^2^ =.102, 95% CI = -0.1594 to 0.02364]. The response criterion, *c*, independent from *d’*, indicates participants’ biases towards responding “old/same”. The mean c for the subjects were [SR: x̄ = -0.047, σ = 0.257; OR: x̄ = -0.148, σ = 0.239] (Figure 5B). Overall, subjects were biased towards responding that faces were remembered as evidenced by negative mean criterion values. However this bias was significantly greater for OR than SR faces [t(21) = 1.733, p = 0.0489. r^2^ = 0.1251, CI = -0.2227 to 0.02023], though the effect was small.

**Figure 5.**
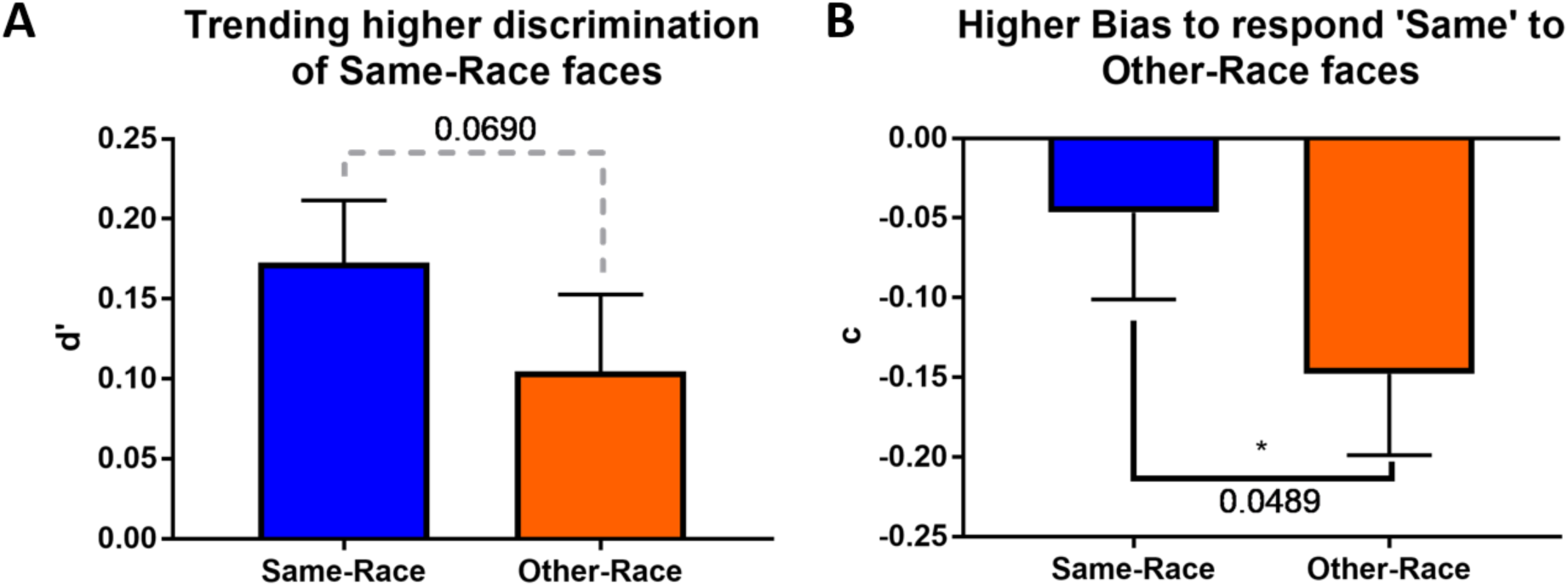
Differences in Recognition Memory for Same- and Other-Race faces. While this task proved difficult for participants, behavioral patterns consistent with the Other-Race Effect still emerged. (A) Sensitivity, *d’*, trended higher for same-race faces [t(21) = 1.542, p = 0.069, r2 =.102, 95% CI = -0.1594 to 0.02364]. (B) Response criterion, *c*, was significantly more liberal for other-race faces [t(21) = 1.733, p = 0.0489. r2 = 0.1251, CI = -0.2227 to 0.02023].

Taken together, these results indicate that this task was difficult – more so than our previously published behavioral variant^27^, but that differences consistent with the canonical Other-Race Effect still emerge. Decreased overall performance on this task may have resulted from an increased time commitment, discomfort, and fatigue associated with studies employed in MRI scanners. In addition, there were several amendments to this task that may have increased difficulty, including requiring indication of left/right face position during encoding as well as the addition of perceptual discrimination trials. Despite this, participants were significantly more likely to answer that they had seen OR faces before, and they demonstrated a trending improved ability to discriminate between old and new SR faces. Furthermore, based on differences in discriminability performance in the (previously published) behavioral version of this task ^27^, the increased difficulty of the scanner task may have impacted SR more than OR recognition. As evidence of this, in the behavioral version, the mean d’ for SR and OR recognition were .3719 and .1484, respectively. In the scanner version of the task there was a 53.59% decrease in same-race discriminability and a smaller 29.45% decrease in other-race discriminability relative to the behavioral study. Moreover, even with the increased difficulty of the scanner version of the task, the subjects’ mean same-race discriminability was 14.02% greater than the mean other-race discriminability in the behavioral task. (Response criterion was not calculated for the behavioral task and is therefore not compared.) This highlights that other-race recognition in less-demanding contexts may still be inferior to same-race recognition in incredibly difficult contexts.

### Neuroimaging

After running through the pipeline described in methods, each of the 16 unique conditions per participant were characterized by global and local efficiency metrics (Equations 1 and 3). Collapsed across conditions, the mean, standard deviation and range of the measures were [Global efficiency: x̄ = 0.5402, σ = 0.0060, range = 0.0476; Local Efficiency: x̄ = 0.4977, σ = 0.0289, range = 0.2144]. Amongst the dataset there was one subject with an outlier condition that was 6.36 standard deviations below the global efficiency mean, and 4.05 standard deviations above the local efficiency mean. This was the only condition present as an outlier for both metrics and was 2.68 and 1.27 standard deviations removed from the next largest outliers for global, and local efficiency, respectively. Furthermore, it had large influence over the relationship between global and local efficiency, inducing a trending correlation which was not present with its removal. [With outlier, r = -0.0909, p = .0885; Without outlier, r= -.0192, p = .7201). Therefore, the extreme outlier was removed, resulting in final descriptive statistics of [Global efficiency: x̄ = 0.5403, σ = 0.0056, range = .03172; Local Efficiency: x̄ = 0.4974, σ = 0.0283, range = 0.1778]. Each of the 16 task conditions corresponds to a distinct phase of the task (study or test), and trial type (target pair or lure pair). Data were sub-grouped based on these conditions to create four distinct categories: A) Study Target, B) Study Lure, C) Test Target, D) Test Lure. Respectively these categories correspond to the conditions under which subjects A) studied faces that would later be repeated, B) studied faces that would later be prompted with similar-looking lure distractors, C) were tested for memory of repeated faces, and D) were tested for ability to correctly reject similar but never-before-seen faces. Within each of these four conditions, data were divided into factors of stimulus race (SR or OR) and accuracy (correct or incorrect). Because each face in the Study Phase had a paired repeat or lure during the Test Phase, we were able to evaluate both brain connectivity during memorization as a function of subsequent performance, and during retrieval. The mean and standard deviations of global and local efficiency for each of these conditions are reported in Table 1.

**Table 1:**
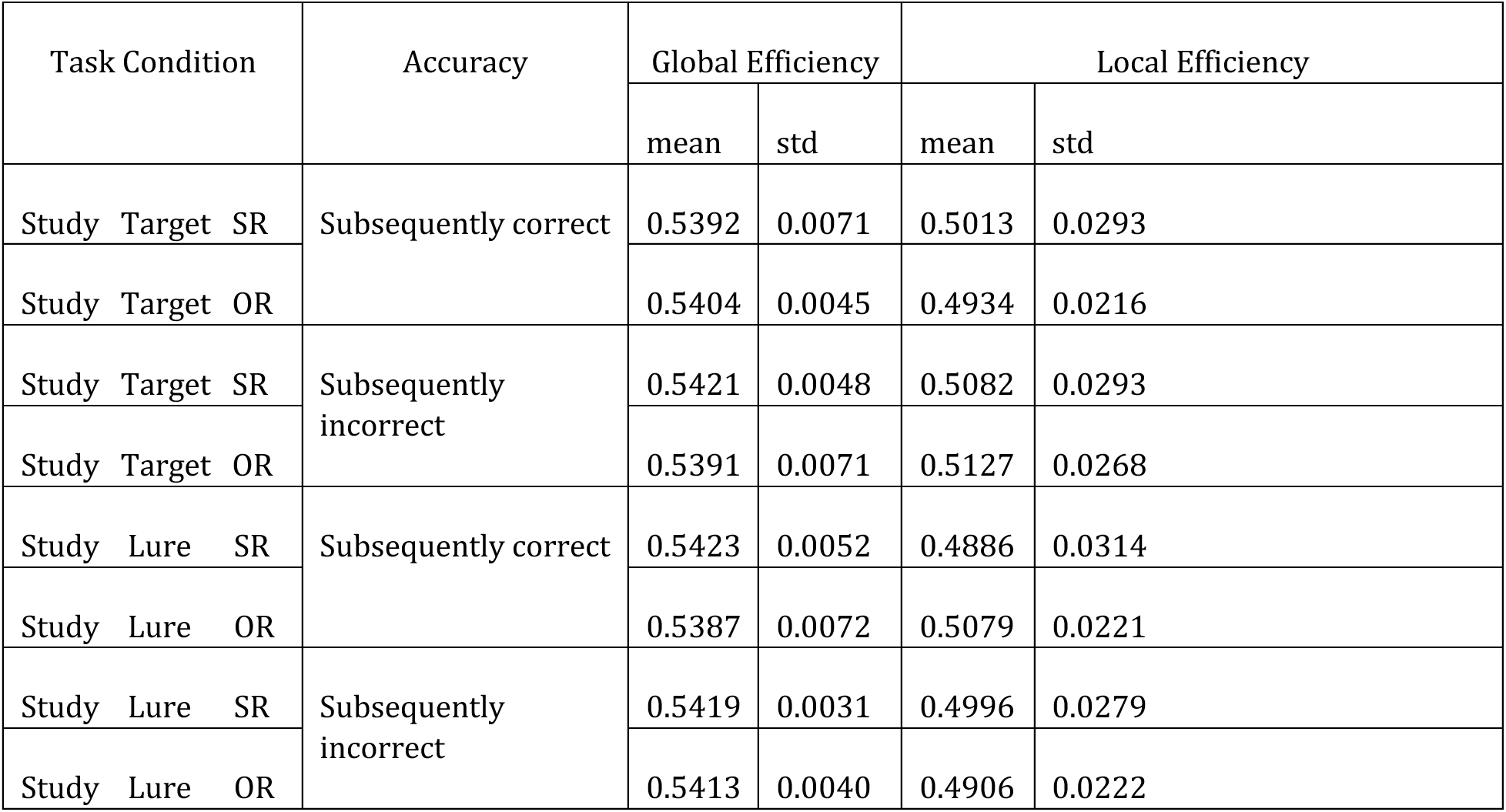
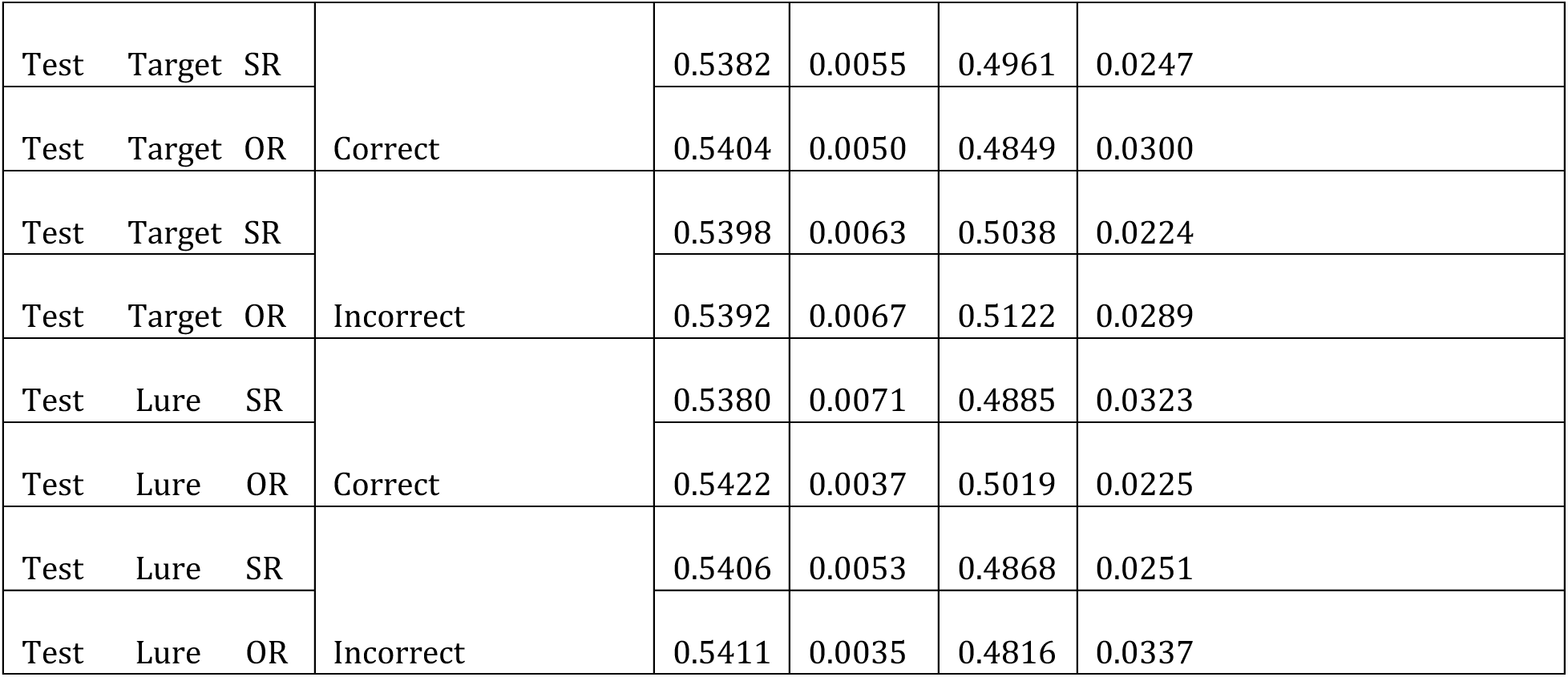
Descriptive Statistics for Global and Local Efficiency During Each Task Condition. Statistics are organized by Task phase (Study or Test), Trial type (Target or Lure), Race (Same-Race or Other-Race), and Accuracy (Correct or Incorrect). For the Study conditions, Target and Lure assignment is based upon whether the face shown was part of a target pair (where the face was later repeated at Test) or a lure pair (where a similar distractor lure was shown at Test). Therefore, Study Accuracy is based upon performance on the corresponding pair during the Test phase.

Within each of the four major categories, a separate regression was run on global and local efficiency data, resulting in 8 models. Specifically, the effects of stimulus race and accuracy on the efficiency metrics were estimated using generalized linear models (GLM) with generalized estimating equations (GEE)^46^. GLMs do not assume normality of dependent variables, and can handle missing values (i.e., the removed outlier), and, when implemented with GEEs, GLMS can handle repeated measures data. We characterized the correlated measures in our data using an exchangeable covariance matrix indicating that all observations per subject were equally correlated to one another. Each regression analysis modeled the linear combination of race, accuracy, and the interaction of race and accuracy on the dependent graph metric (Equation 6).

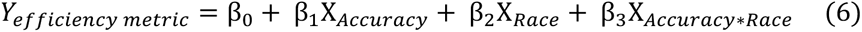

Because the ORE is behaviorally characterized by differences in performance relative to the race of faces, we were specifically interested in identifying whether efficiency of information processing in the brain is modulated by race and accuracy. We therefore directed our focus to the interaction of race and accuracy (β_3_ X_Accuracy*Race_). Given the small sample size (22 observations per factor), we ran permutation analysis for each model to reduce the likelihood that results could be due to chance. For each observed dataset, 10,000 randomized datasets were generated where each of the four corresponding condition labels were shuffled within each subject. The 10,000 null datasets were then modeled with GLM/GEEs using the same pipeline described above. To evaluate significance, we calculated the proportion of times that the permutation analysis estimated larger interaction test statistics, z, than those produced when modeling the true data. All 8 uncorrected p-values were then subjected to the Benjamini-Hochberg procedure to control the false discovery rate (FDR). Table 2 includes a summary of estimates for the interaction terms across conditions, including beta coefficients, z-scores, 95% confidence intervals, p_uncorrected_ and p_adjusted._

**Table 2:** Estimates for the Interaction of Race and Accuracy Across All Conditions. The columns for β, z, and 95% Confidence Interval (CI) are results from the original models. In generalized linear models (GLM) with generalized estimating equations (GEE), the z-score statistics are calculated by dividing the beta estimates by robust standard errors, which are larger and more conservative than traditional standard errors. p_uncorrected_ was calculated by permutation analysis, while p_adjusted_ reflects false discovery rate (FDR) corrections after the Benjamini-Hochberg procedure.

| Phase | Trial Type | Graph Metric | $\beta$ | $z$ | 95% CI | $p_{\text{uncorrected}}$ | $p_{\text{adjusted}}$ |
| --- | --- | --- | --- | --- | --- | --- | --- |
| Study | Target, 1<sup>st</sup> Presentation | Global efficiency | 0.0042 | 1.674 | [-0.001, 0.009] | 0.0586 | 0.09376 |
|  |  | Local efficiency | -0.0123 | -1.332 | [-0.030, 0.006] | 0.102 | 0.1099 |
|  | Lure, 1<sup>st</sup> presentation | Global efficiency | -0.0029 | -1.468 | [-0.007, 0.001] | 0.0869 | 0.1099 |
|  |  | Local efficiency | 0.0282 | 3.088 | [ 0.010, 0.046] | 0.0028 | 0.0224 |
| Test | Target Repeat | Global efficiency | 0.0028 | 1.279 | [-0.002, 0.007] | 0.1099 | 0.1099 |
|  |  | Local efficiency | -0.0196 | -2.026 | [-0.039, -0.001] | 0.029 | 0.07733 |
|  | Lure Distractor | Global efficiency | 0.0037 | 1.800 | [-0.000, 0.008] | 0.0457 | 0.0914 |
|  |  | Local efficiency | 0.0186 | 2.402 | [ 0.003, 0.034] | 0.0154 | 0.0616 |

As indicated in Table 2, four of the eight models had significant interaction terms prior to FDR correction (p_uncorrected_ < .05). After adjustments, only the Study Lure condition maintained a significant interaction effect on network local efficiency [p_uncorrected_ = .0028, p_adjusted_ = .0224, z = 3.088, β = .0282, SE_robust_ = .009, CI = [0.010, 0.046]]. While the other three effects remained trending, they corresponded to retrieval conditions, and have much reduced test statistics [Test Target_local_, z = -2.026; Test Lure_global_, z = 1.800; Test Lure_local_, z = 2.402]. Given this, the remainder of our analysis is post-hoc, and focuses on the Study Lure condition.

Figure 6A depicts the modulation of local efficiency during encoding by both race and accuracy factors. For faces that were encoded well enough to avoid false alarms at retrieval, mean local efficiency was greater for OR faces [x̄_SR_ = .4886; x̄_OR_ = 0.5079]. Meanwhile, the opposite trend is demonstrated for encoding resulting in false alarms, with greater local efficiency for SR faces [x̄_SR_ = .4996; x̄_OR_ = 0.4906]. (Refer to Table 1 for descriptive statistics for all conditions.) This cross-over interaction suggests that SR and OR face recognition are supported by divergent segregation properties of brain connectivity and information exchange. Only encoding of OR faces was more successful when the brain network was organized to enhance fault tolerance via localized clustering of nodes preserving redundant connections.

**Figure 6:**
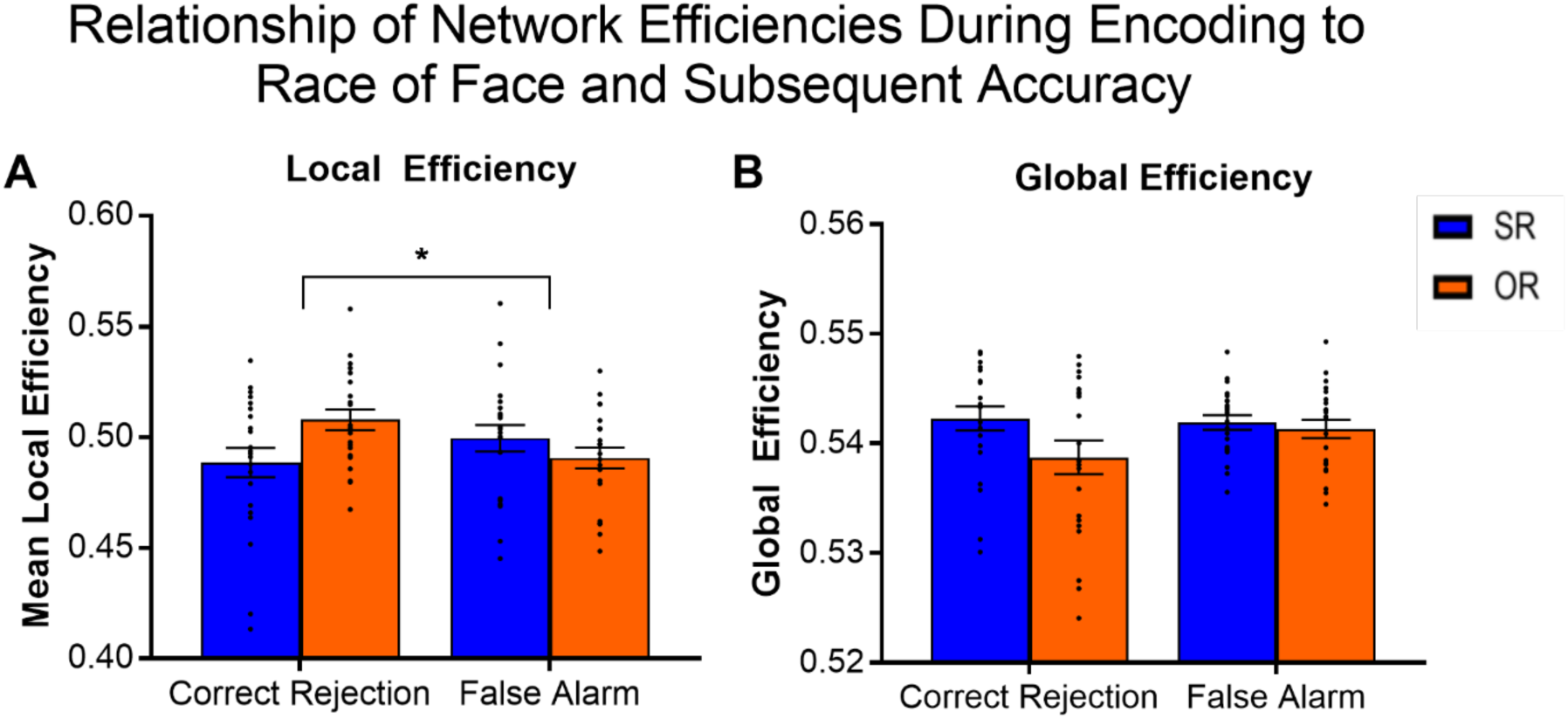
Local and Global Efficiency during Encoding. Network efficiency in the brain during encoding of faces that later resulted in correct rejections or false alarms. (A) There was a robust interaction of race and subsequent accuracy on mean local efficiency [p_uncorrected_ = .0028, p_adjusted_ = .0224, z = 3.088, β = .0282, SE_robust_ = .009, CI = [0.010, 0.046]]. B) A post-hoc analysis found a main effect of race on global efficiency [z = 2.067, β = 0.0036, SE_robust_ = .002, p = 0.039, CI = [0.000, 0.007]].

In contrast, encoding of SR faces was more successful when local organization was more distributed, with sparser interconnectivity between close functionally connected nodes. Given that all networks were kept comparable by maintaining equal edge counts, this poses the question of how edges were distributed, and whether they might be supporting a more globally integrated network. Though permutation analysis established there was no interaction of race and accuracy on global efficiency during lure encoding, we tested a post-hoc hypothesis that SR face encoding might be characterized by a more globally integrated network in this condition (Figure 6B). We reran the regression analysis on the study lure global efficiency data and extracted the β estimates for the main effect of race, rather than the interaction. The results suggest that race had a significant impact on global efficiency during the study lure condition [z = 2.067, β = 0.0036, SE_robust_ = .002, p = 0.039, CI = [0.000, 0.007]]. Figure 6B displays this main effect, where global efficiency is enhanced during encoding of SR relative to OR faces. [x̄_SR_Corr_ = 0.5423; x̄_OR_Corr_ = 0.5387; x̄_SR_Incorr_ = 0.5419; x̄_OR_Incorr_ = 0.5413]. Despite this being a main effect, it appears more strongly driven by the correct rejection condition.

Thus far, all data reported have been whole-brain network-level metrics. Recall, to calculate local efficiency, nodal subgraph efficiencies were averaged together for the entire network. Because each node has its own local efficiency, we additionally tested whether the observed pattern of greater network segregation during accurate face encoding of OR faces was uniform across discrete sub-networks in the brain, or whether any network in particular drove the overall higher mean. Research has shown that at rest, the brain demonstrates segregated functional connectivity that results in separate modules^21^. The integrity of these modules, often called intrinsic connectivity networks (ICNs), are associated with varying cognitive functions, such as visual perception, bottom-up and top-down attention, cognitive control, etc^47^. It was therefore of interest to test whether networks associated with certain perceptual and attentional capacities might be differentially engaged for same- and other-race face recognition.

To test this, each node was assigned to an ICN using a preexisting mapping between the Brainnetome atlas and Yeo ICN parcellations^36,47^. The ICNs included in this parcellation are the Dorsal Attention Network (DAN), Default Mode Network (DMN), Frontoparietal Network (FPN), Limbic Network, Somatomotor (SOM) Network, Ventral Attention Network (VAN) and Visual Network. In the mapping, 207 out of the full sample of 243 nodes corresponded to one of the 7 ICNs. The remaining 36 nodes were removed from the analysis, 33 of which were subcortical and not included in the Yeo Parcellation. The final 3 nodes that were removed corresponded to portions of the cingulate gyrus and insula. The final number of nodes per ICN are described in Table 3.

**Table 3.**
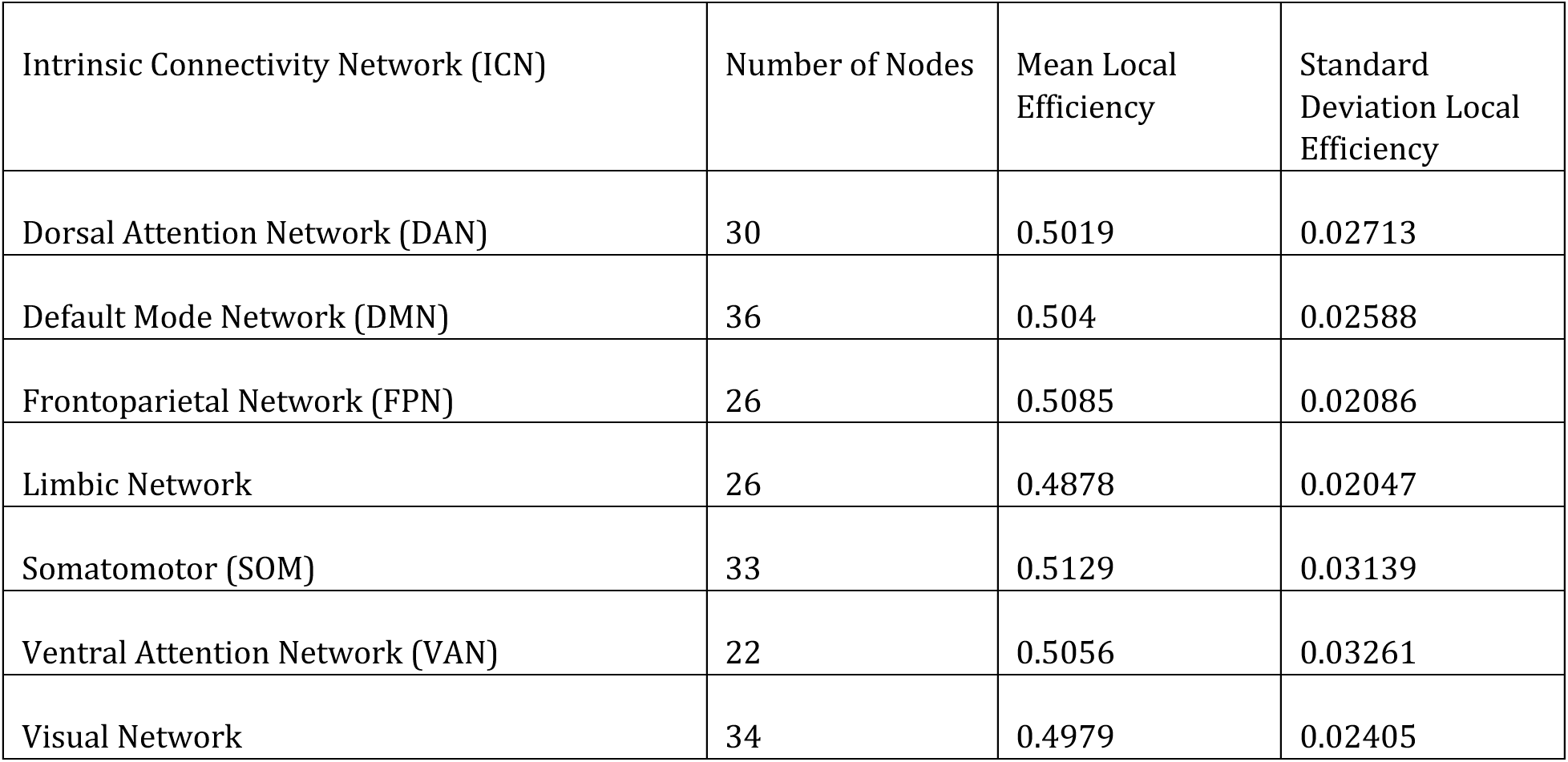
Descriptive Statistics for Local Efficiency Across Intrinsic Connectivity Networks. Of the 246 nodes in the Brainnetome brain networks, 207 mapped onto known intrinsic connectivity networks (ICNs) from the Yeo parcellation. The number of nodes included in each network is shown above, as well as the mean and standard deviation of local efficiency within each network.

Next, ICN-level mean local efficiencies were computed for each subject, by averaging together the local efficiency of nodes within each ICN across the four study lure conditions. It should be noted that mean local efficiency of an ICN reflects the mean efficiency with which its nodes’ direct neighbors are coupled, independent of the neighbors’ ICN affiliation. That is, mean efficiency of an ICN should be interpreted as the overall fault tolerance of the subgraphs of the ICN nodes. This calculation does *not* comment on the efficiency with which nodes within a shared ICN are coupled. Mean efficiency was calculated with conditions collapsed (Table 3) as well as for each separate condition, plotted in Figure 7 Figure 8 to emphasize differences across the race and accuracy factors.

**Figure 7.**
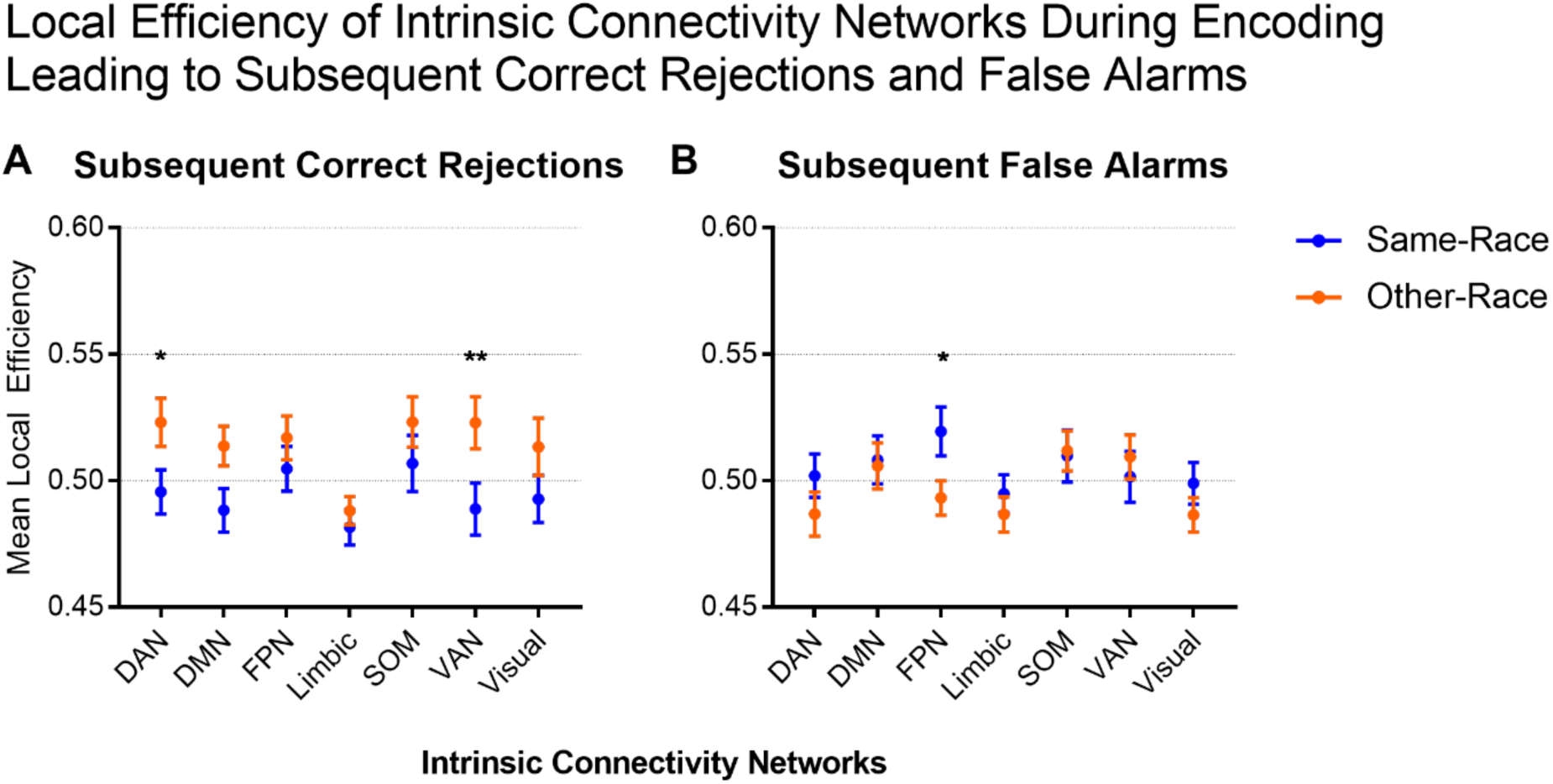
Facial Race Impacts on Local Efficiency of Intrinsic Connectivity nodes during Accurate and Inaccurate Encoding. A) Across intrinsic networks (ICNs), encoding promoting subsequent correct rejections was associated with greater local efficiency for OR faces. After an ANOVA and multiple comparison correction, DAN and VAN efficiency remain statistically different across race. B) There is less consistency in encoding promoting subsequent false alarms. SR tends to, but does not always demonstrate higher local efficiency. After multiple-comparison correction, the FPN efficiency remains significantly different across race. Abbreviations: DAN - Dorsal Attention Network; DMN - Default Mode Network; FPN - Frontoparietal Network; SOM -Somatomotor (SOM) Network; VAN - Ventral Attention Network.

**Figure 8.**
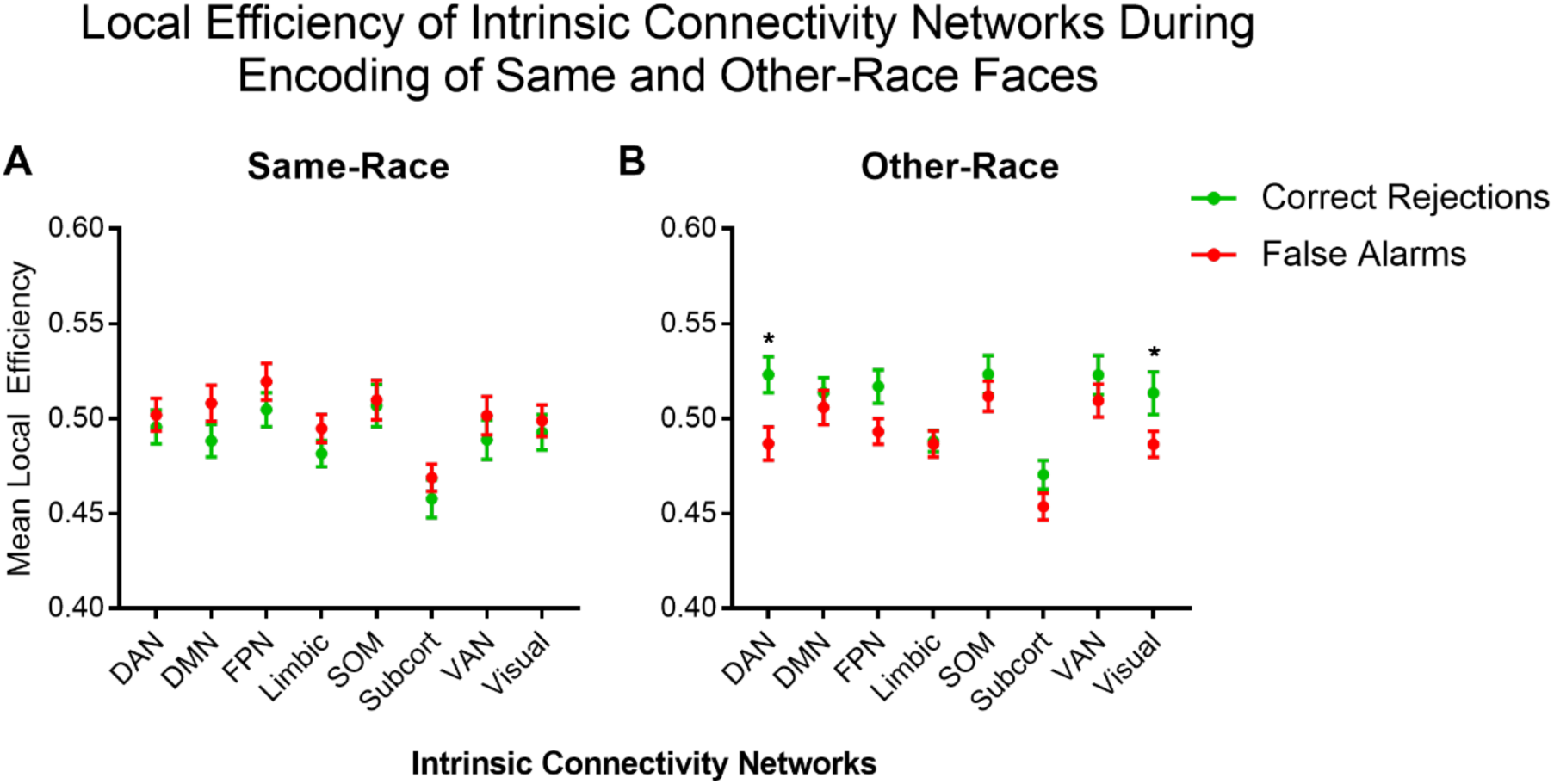
Accuracy Differences in Local Efficiency of Intrinsic Connectivity nodes during Same and Other-Race Face Encoding. (A) During encoding of SR faces, local efficiency of ICNS does not appear to be significantly associated with subsequent accuracy. (B) Meanwhile, there is a relationship between accuracy and local efficiency in the OR condition, with higher efficiency during encoding of subsequently accurate trials. After multiple-comparison correction, the DAN and Visual networks maintain significantly greater efficiencies for the correct rejection relative to false alarm condition. Abbreviations: DAN - Dorsal Attention Network; DMN - Default Mode Network; FPN - Frontoparietal Network; SOM -Somatomotor (SOM) Network; VAN - Ventral Attention Network.

Figure 7A highlights that the increase in local efficiency during accurate encoding of OR relative to SR faces was replicated across intrinsic connectivity networks. Meanwhile, efficiency during inaccurate encoding (Figure 7B) across networks is not consistently different between same- and other-race conditions. Still, the cross-over interaction between race and accuracy observed in the whole-brain networks (Figure 6A) is reproduced in a 3-way repeated measures ANOVA with factors of race, accuracy, and ICN. We find both main effects of ICN [F(6,588) = 3.33, p = .0031, η2 = .03] and an interaction between race and accuracy [F(1,588) =17.56, p<.0001, η2 = .03] but no three-way interaction [F(6,588) = 0.3758, p = .89, η2 = 0]. We next tested whether specific subnetworks were driving whole-network differences, since certain ICNs display non-overlapping standard error of means across the race conditions (Figure 7A,B). To do this, two-way repeated measures ANOVAS with factors of race and ICN were applied separately on the correct rejection and false alarm conditions. Post-hoc Sidak correction was applied to test which subnetworks showed the greatest deviation in local efficiency across race. For subsequent correct rejections there were main effects of race [F(1,21) = 7.72, p =.011, η2 = .05] and ICN [F(6,126) = 2.92, p =.011, η2 = .04]. After correcting for multiple comparisons across the eight networks, the increased local efficiency in OR relative to SR encoding remained significant in the DAN [SR x̄ = .50, OR x̄ = .52, p = .04, 95% CI = [- .05 to 0]] and VAN [SR x̄ = .49, OR x̄ = .53, p = 0.005, 95% CI = [-0.06 to -0.007]]. For the same ANOVA applied to the subsequent false-alarm condition, there was no effect of race, but the effect of ICN remained [F(6,126) = 2.43, p =.03, η2 = .03]. The lack of effect of race in the inaccurate condition suggests that the differences observed in local efficiency across race in the accurate condition promote successful same- and other-race memory. Despite no overall main effect of race, the FPN demonstrated a significant difference with respect to race after Sidak correction [SR x̄ = .52, OR x̄ = .49, p = 0.02, 95% CI = [0 to 0.05]], with greater SR relative to OR efficiency associated with subsequent false alarms. Note, the directionality of effects on DAN, VAN, and FPN mirror the overall significant effects on the whole-brain networks (Figure 6).

We additionally tested whether specific ICNs drove differences within race across accuracy conditions. Figure 8 displays local efficiency across accurate and inaccurate encoding conditions for SR (A), and OR (B) faces. Because the standard error of mean bars overlap for all networks in the SR condition, no follow-up tests were run. Because network efficiency across accuracy appears more distinct in the other-race condition, a two-way repeated measures ANOVA was applied with factors of accuracy and ICN followed by a post-hoc Sidak correction for multiple comparisons. This resulted in main effects of accuracy [F(1,21) = 8.21, p = .009, η2 = .04] and ICN [F(6,126) = 3.09, p =.007, η2 = .05], with accurate trials generally being associated with higher local efficiency across networks. These differences in efficiency across ICNS survived corrections for DAN [CR x̄ = .52, FA x̄ = .49, p = .002, 95 % CI = [0.01 to 0.06]] and the Visual Network [CR x̄ = .51, FA x̄ = .49, p = .045, 95 % CI = [0 to 0.05]].

Considering whole-brain and ICN-level efficiencies together, the result of this study suggests that functional reorganization of the brain during encoding influences subsequent capacity to correctly reject face lures. Distinct brain network topologies supported behavior relative to the race of presented faces. Successful same-race encoding was related to a more integrated network infrastructure, while accurate other-race encoding was associated with more network segregation. Greater segregation during successful other-race encoding was widely distributed across functional subnetworks of the brain. Furthermore, local efficiency during encoding across ICNS provided an accuracy advantage in other-race faces, but not same-race faces.

## Discussion

The present study demonstrates successful same-race (SR) and other-race (OR) face recognition are supported by differing functional brain network topologies. Connectivity in the brain during encoding was greatly modulated by the race of presented faces. This suggests that the context of race alters the behavioral adaptivity of network configurations. OR face encoding promoting correct rejections was subserved by a more locally efficient network, while reduced local efficiency was associated with false identifications. In contrast, SR recognition was associated with the opposite pattern: reduced local efficiency during encoding supported later correct rejections, while greater local efficiency was related to false alarms. The connection between SR accuracy and reduced segregation was further underscored by greater global efficiency during encoding of SR relative to OR faces. Together these data suggest that a more functionally integrated network is important for SR correct rejections, while a more segregated network is optimal for OR correct rejections.

These results imply that there is not a one-size fits all network topology for successful face recognition. Across study and retrieval conditions tested, we found four of eight conditions demonstrated interactions between race and accuracy, though only the study lure condition survived post-hoc correction. While research often associates integration with memory accuracy ^21,22,48^, there are cases, like the OR context, where more segregated networks support success. One such study found that a working memory task was only initially supported by an integrated brain network in participants, but with training, improvements in performance were coupled with more modular network configurations^23^. These findings suggest segregated networks may still play a role where performance advantages are concerned. In contrast, SR recognition was subserved by globally integrated networks, which is more consistent with literature on network configurations supporting accurate memory formation.

While to our knowledge there have been no developmental analyses of the ORE from a graph theoretical perspective, several studies have used graphs to understand general facial recognition across development^49,50^. One cross-sectional study found that across development there is, in fact, an increase in integration between face-preferential regions in visual and limbic networks. Our results align with these, while suggesting that this integration is widespread beyond face-selective regions and is specific to SR faces with which we have increased experience with. This globally efficient configuration may reflect an ease with which information about SR faces can be propagated across the brain. The counter-effect we observed of increased segregation during accurate OR face discrimination may reflect a compensatory configuration of specialized modules supporting interaction with faces deviating from those that we have been expertly trained on over our lifetimes.

We additionally found differences in how intrinsic connectivity networks processed SR and OR face information. Regions in the ventral attention network (VAN) showed the largest increase in local efficiency for successful OR relative to SR encoding. However, all subnetworks demonstrated higher local efficiency to OR faces, suggesting that widespread functional reorganization promoting network redundancy may be especially important for OR encoding. Furthermore, brain regions in the visual and dorsal attention networks (DAN) were more locally efficient during accurate than inaccurate OR encoding, while local efficiency across networks during SR encoding seemed to have less bearing on overall performance.

It is intriguing that the subnetwork-level analysis implicates that network segregation involving visual, top-down (DAN) and bottom up (VAN) regions was specifically important during encoding of OR faces. Visual regions have been most greatly explored in the context of the ORE (Fusiform Face Area, Occipital Face Area), however nodes within the VAN and DAN are also included in the core and extended face processing networks^51^. For instance, the posterior superior temporal sulcus in the VAN is considered a core face region involved in directing attention during face encoding and has demonstrated differential activity relative to the race of face stimuli^52^. Other VAN regions like the insula and cingulate gyrus are involved in salient emotional face processing^53,54^. In addition, the VAN is globally involved in stimulus-driven attentional control and may be recruited to orient attention divergently across SR and OR conditions. Meanwhile, DAN nodes like the intraparietal sulcus help direct eye gaze and the middle temporal gyrus is implicated in familiar face processing^52^. The DAN is also involved in task-driven attentional control, biasing responses from lower-level visual regions. All of these cognitive functions are vital for face recognition, so it is possible that specialized and redundantly organized modules may form between nodes included in these networks, enabling more successful encoding of OR faces.

The fact that no major differences were found in local efficiency across networks associated with accurate and inaccurate SR encoding is noteworthy. It is possible that perceptual expertise reduces the need to encode information as redundantly, since a more integrated network could compensate for reduced local fault tolerance. This integration could involve highly central nodes/hubs which could computationally lead to reduced local efficiency due to an inability for networks to be resilient to their removal. However, the extent to which several hubs could influence mean efficiencies in entire subnetworks is unclear. Future research could test whether certain hubs exist during SR but not OR recognition, and whether this impacts mean local efficiency of intrinsic networks.

The intrinsic connectivity level results found here are bolstered by findings from a study testing the importance of top-down attentional and control networks in shaping the ORE^16^. Differences in activity in the intraparietal sulcus (associated with DAN and Fronto-parietal cognitive control networks) were particularly associated with failures to remember OR faces. Furthermore, functional connectivity between the right fusiform cortex and intraparietal sulus regions in the DAN were significantly greater for SR than OR faces. It is important to note that the study design, analyses and results diverge from ours, in that success was defined as target recognition (as opposed to lure correct rejection), and analyses were not explored from a graph theoretical perspective. Still, both studies find that there is altered behaviorally relevant recruitment of regions of the DAN during encoding of SR and OR faces. Our more exploratory study contributes the novel observation that altered topology in the VAN and visual intrinsic connectivity networks may also play a role in the ORE. Future more targeted studies may be designed to focus on whether the interactions (or lack thereof) of these networks might relate to the emergence of SR and OR recognition disparities.

This study’s key findings pertain specifically to encoding that later subserved successful mnemonic discrimination ^27^ -- the ability to distinguish between similar but distinct memories or experiences/ Although we noted three potential interactions between race and accuracy in brain connectivity during retrieval, these were not further analyzed as they were only trending after multiple comparison corrections. We further did not find that network-based interactions between race and accuracy was associated with successful target recognition. As target recognition memory deficits are as central to the ORE as mnemonic discrimination, our results on mnemonic discrimination represent only a part of a larger phenomenon.

Future network neuroscientific investigations of the ORE will benefit from the following considerations: While the simultaneous inclusion of encoding, retrieval, target, and lure conditions was novel, it greatly reduced our power to detect significant effects given the need for stringent post-hoc corrections. Subsequent studies should consider more targeted hypotheses and experimental designs. In addition, the large number of conditions increased the length of this study, likely contributing to it being overly difficult for participants. Despite low performance, a highly conservative analysis found that differences in brain network topologies across encoding lure contexts were likely not due to chance. Still, future studies can be improved by shortening the task design and increasing physiognomic differences between faces in lure pairs to reduce task complexity. Furthermore, while there are a variety of ways to analyze task-based connectivity in the brain, no large systematic study has looked at differences in resulting graph topologies based on choice of functional connectivity method. Because graph theoretical analysis may be sensitive to selection of brain atlas, preprocessing methods, and functional connectivity analysis, it is important to study the network-based correlates of the Other-Race Effect across a variety of paradigms and methods to ensure that the findings reported here are reliable and reproducible.

In conclusion, participants performed a mnemonic discrimination paradigm in the MRI scanner, allowing us to evaluate the role of face recognition memory processes on the functional network organization of the brain. To test this, we computed network topology metrics from graph representations of functional connectivity during multiple recognition conditions for SR and OR faces. Our findings suggest that SR and OR face encoding is supported by distinct network topologies. The ability to not mistakenly recognize lure faces at retrieval was supported by redundantly organized modules during encoding of OR faces, and by a more integrated network during encoding of SR faces. The relatively greater importance of network segregation for OR faces was supported by greater efficiency in visual, bottom-up, and top-down attentional networks during accurate compared to inefficient encoding. As the first (to our knowledge) graph theoretical analysis specific to the ORE, these results demonstrate that face recognition is modulated by the relationship the viewer has with the race of the face they are processing, causing large-scale whole-brain differences in functional network communication. These findings should serve to motivate research beyond face-preferential regions to understand how systems-wide connectivity differences promote the ORE.

